# A stepwise route to domesticate rice by controlling seed shattering and panicle shape

**DOI:** 10.1101/2021.12.02.470680

**Authors:** Ryo Ishikawa, Cristina Cobo Castillo, Than Myint Htun, Koji Numaguchi, Kazuya Inoue, Yumi Oka, Miki Ogasawara, Shohei Sugiyama, Natsumi Takama, Chhourn Orn, Chizuru Inoue, Ken-ichi Nonomura, Robin Allaby, Dorian Q Fuller, Takashige Ishii

## Abstract

Asian rice (*Oryza sativa* L.) is consumed by more than half of the world’s population. Despite its global importance, the process of early rice domestication remains unclear. During domestication, wild rice (*O. rufipogon* Griff.) acquired non-seed-shattering behaviour, allowing humans to increase grain yield. Previous studies argued that the *sh4* mutation triggered a reduction in seed shattering during rice domestication; but our experiments using wild introgression lines of *O. rufipogon* show that the domesticated *sh4* allele alone is insufficient for shattering loss. Here, we identified the interaction between three key mutations associated with the interruption of abscission layer formation and panicle architecture that were causal in early rice domestication. An interruption of abscission layer formation requires both *sh4* and *qSH3* mutations, presenting an apparent barrier to the selection of shattering loss. We identified a causal single-nucleotide polymorphism at *qSH3* within the seed-shattering gene *OsSh1*, which is conserved in *indica* and *japonica* subspecies but absent in the *circum*-aus group of rice. Through harvest experiments, we demonstrated that seed shattering alone did not significantly impact yield; rather yield increases were observed with closed panicle formation controlled by *SPR3*, which is further augmented by the integration of *sh4* and *qSH3* alleles. Complementary manipulation of seed shattering and panicle shape result in a mechanically stable panicle structure. We propose a stepwise route for the earliest phase of rice domestication, wherein selection of visible *SPR3*-controlled closed panicle morphology was instrumental in the sequential recruitment of *sh4* and *qSH3*, which led to the loss of shattering.

**Significance Statement:** Rice is one of the most important crops worldwide. Loss of seed shattering in domesticated rice, previously attributed to single mutations such as those in *sh4*, is considered the principal genetic change which resulted in yield increases. However, we show that *sh4* alone is insufficient and other genes, such as *qSH3*, are required to cause abscission layer disruption. The evolution of non-seed-shattering therefore required multiple mutations. Furthermore, shattering loss in genetic backgrounds of wild rice does not correspondingly increase yields. We have identified an interaction in which a second trait, closed panicle formation controlled by *SPR3*, that both increases the yield and facilitates recruitment of *sh4* and *qSH3*, which synergistically augment yield, leading to a stepwise model for rice domestication.

## Introduction

The selection of naturally occurring variations in wild plants that provide useful agronomic traits is an essential step in crop domestication. These traits are often related to yield, such as seed number, seed size, and a loss of seed dispersal, as well as ease of cultivation, including plant architecture, seed dormancy, and photoperiod sensitivity. Plants with these traits provided the necessary impetus for humans to shift from hunting and gathering to cultivation. Among several domestication-related traits, the suppression of seed shattering is considered the most important genetic change that allowed humans to increase grain yield and differentiated domesticated from wild plants (1, 2). However, the role of visible morphological changes in facilitating domestication is debated.

Rice (*Oryza sativa* L.) is consumed by almost half of the world’s population and is particularly important in Asia. During its domestication, wild rice (*O. rufipogon* Griff.) was transformed by acquiring non-seed-shattering behaviour. Investigations on prehistoric rice spikelet bases from archaeological sites demonstrate that suppression of seed shattering leaves a phenotypic trace that can be observed through the examination of the abscission layer (3). Archaeological rice spikelet bases with rough and deeply torn scars in the spikelet base (rachilla) indicate that humans had to actively detach the spikelets from the pedicels by threshing. More than a decade ago, two seed-shattering loci, *sh4* and *qSH1*, were identified as essential to rice domestication (4, 5). The mutation at *sh4* was largely viewed as the principal genetic change that led to rice domestication. Quantitative trait locus (QTL) analysis identified *sh4* as the locus responsible for the difference in seed-shattering behaviour between wild rice *O. nivara* (an annual form of *O. rufipogon*) and *O. sativa* ssp. *indica*. A single-nucleotide polymorphism (SNP) at *sh4* was found to affect the function of the Trihelix transcription factor due to an amino acid change in its DNA-binding domain, resulting in suppression of seed shattering (4; Table S1). *qSH1* was also detected as the QTL largely responsible for the difference in seed-shattering degree between the rice cultivars, *O. sativa* ssp. *japonica* and *O. sativa* ssp. *indica*. A SNP of *qSH1* resulted in suppression of seed shattering in *O. sativa* ssp. *japonica* cv. Nipponbare by downregulating expression of the downstream gene *OsRIL1* encoding a BEL1-type homeobox family protein (5; Table S1). Since *sh4* and *qSH1* display large phenotypic variances, and rice cultivars with functional alleles of *sh4* or *qSH1* promote seed shattering, they are recognised as crucial to rice domestication.

To better understand if the domestication process of Asian rice was promoted mainly by a single domestication gene, such as *sh4*, and also to see how *qSH1* and *sh4* are involved in the loss of seed shattering in wild rice, we previously developed the introgression lines (ILs) named IL(*qSH1*-N) and IL(*sh4*-N). These ILs contain small chromosomal segments of domesticated-type from *O. sativa* Nipponbare at the *qSH1* and *sh4* loci, respectively, in the genetic background of wild rice *O. rufipogon* W630 (6). Complete seed shattering was still observed in these lines (6–8), indicating that neither of the single mutations at *qSH1* and *sh4* on its own could account for non-shattering leading to rice domestication. As the *sh4* mutation is commonly observed in cultivated rice, while the *qSH1* mutation is only found in some *japonica* cultivars (5), additional mutation(s) together with *sh4* must have played a role in reducing seed shattering during the early stages of rice domestication. Such quantitatively inherited mutations reducing shattering behaviour are an apparent barrier to selection of shattering loss which does not occur on the basis of a simple recessive mutation. Thus, it is plausible that the domestication process in rice is more complicated than the previously proposed single domestication allele model.

A change in panicle structure from open to closed may have played a crucial role in mitigating seed-shattering behaviour prior to the selection of non-shattering rice (7). *SPR3* is responsible for closed panicle formation and acts as a cis-regulatory element controlling expression of the downstream gene, *OsLG1*, which encodes a *SQUAMOSA* promoter-binding protein (7). A closed panicle structure encases the mature seeds retained in the upper part of the panicle with the long awns of the lower immature seeds, potentially making these plants more attractive to gatherers. By choosing rice plants with the *SPR3* genetic mutation, gatherers would have increased their collection rate, particularly if the plants with closed panicles also exhibited suppressed seed shattering. In addition, selection of the closed panicle formation also brought about increased self-pollination rates. Therefore, this trait may have potentially contributed to the accumulation and fixation of mutations, leading to the evolution of non-seed-shattering plants thereafter.

In this study, we conducted a genetic analysis of *qSH3*, an additional locus involved in the loss of seed shattering together with *sh4*, and identified the causal mutation selected during rice domestication. The frequency of the *qSH3* alleles in rice cultivars and their effects on the loss of seed shattering were also evaluated. We further conducted seed gathering experiments, which showed how slight genetic changes can significantly impact yields. Finally, we assessed the potential role of closed panicle formation on reducing seed shattering and the relationship between these two distinct traits, seed shattering and closed panicle formation, based on structural mechanic analysis to better understand the selection process on a loss of seed shattering in rice.

## Results and discussion

### Identification of the causal mutation at *qSH3*, a locus involved in the loss of seed shattering during rice domestication

To identify additional genomic regions involved in reducing seed shattering, we previously produced an IL with chromosomal segments of *O. sativa* Nipponbare at both *sh4* and *qSH1* in the genetic background of wild rice, *O. rufipogon* W630 (8; Fig. S1). IL(*qSH1*-N, *sh4*-N), which contains small chromosomal segments from *O. sativa* Nipponbare at both *qSH1* and *sh4* loci, was crossed with Nipponbare, and their F_2_ population was subjected to QTL analysis to determine the degree of seed shattering, and the locus *qSH3* was identified (8, 9). High-resolution linkage analysis using a mapping population from a cross between IL(*qSH1*-N, *sh4*-N) and Nipponbare (10; Fig. S1) identified part of a previously known seed-shattering gene, *OsSh1* (*Os03g0650000*), a homolog of *Sh1* controlling abscission layer formation in sorghum (11; Fig. S2–S4, Table S1). The function of *OsSh1* in seed shattering was studied using artificially induced rice materials showing that the null mutations caused a complete loss of seed shattering (11, 12), but the causal mutation selected in rice domestication was not determined. No significant difference was detected in *qSH3* expression levels in the spikelet base between Nipponbare and W630 (Fig. S4), suggesting that the *qSH3* expression level may not be a cause of differences in the seed-shattering degree. Among the seven polymorphisms in the region (Fig. S4 and Table S2), only one SNP of C in W630 and T in Nipponbare, namely SNP-70, was located in the coding region (exon1) of *qSH3*, causing an amino acid substitution from leucine in W630 to phenylalanine in Nipponbare (8; Fig. S5, Table S1). To test whether SNP-70 is associated with the degree of seed shattering, we performed a transformation test in the Nipponbare genetic background capable of transgenic experiments using two constructs that differed only at the SNP-70 position. In the preliminary experiment using Nipponbare ILs without transformation (Fig. 1A), IL(*qSH3*-W), carrying the W630 allele at *qSH3* in the Nipponbare genetic background, showed a non-seed-shattering behaviour similar to that of Nipponbare, whereas IL(*qSH1*-W, *qSH3*-W) displayed a significantly higher seed-shattering degree than that of IL(*qSH1*-W) based on the breaking tensile strength (BTS) value (Fig. 1B and Fig. S6). These findings indicated that the seed-shattering effect of *qSH3* was considerably enhanced by a functional allele at *qSH1* (Fig. S7). Therefore, we introduced two types of constructs carrying the *qSH3* cDNA sequence of W630 (*qSH3*^*W630*^) and Nipponbare (*qSH3*^*Npb*^) into IL(*qSH1*-W) (Fig. 1C). The transgenic plants were expected to express both endogenous *qSH3* of Nipponbare in IL(*qSH1*-W) and *qSH3*^*W630*^ or *qSH3*^*Npb*^ as the transgene (Fig. S7). Therefore, we screened transformants expressing transgenes using the derived cleaved amplified polymorphic sequences (dCAPS) assay targeting a SNP at the 5’ UTR, encoded by the W630 promoter region (Fig. S7–S9). As a result, transformants with the wild-type *qSH3*^*W630*^ showed enhanced seed shattering, compared with the control IL(*qSH1*-W), whereas those with domesticated-type *qSH3*^*Npb*^ did not shatter (Fig. 1D, Fig. S7), confirming that SNP-70 is the causal mutation for reduced seed shattering.

**Fig. 1.**
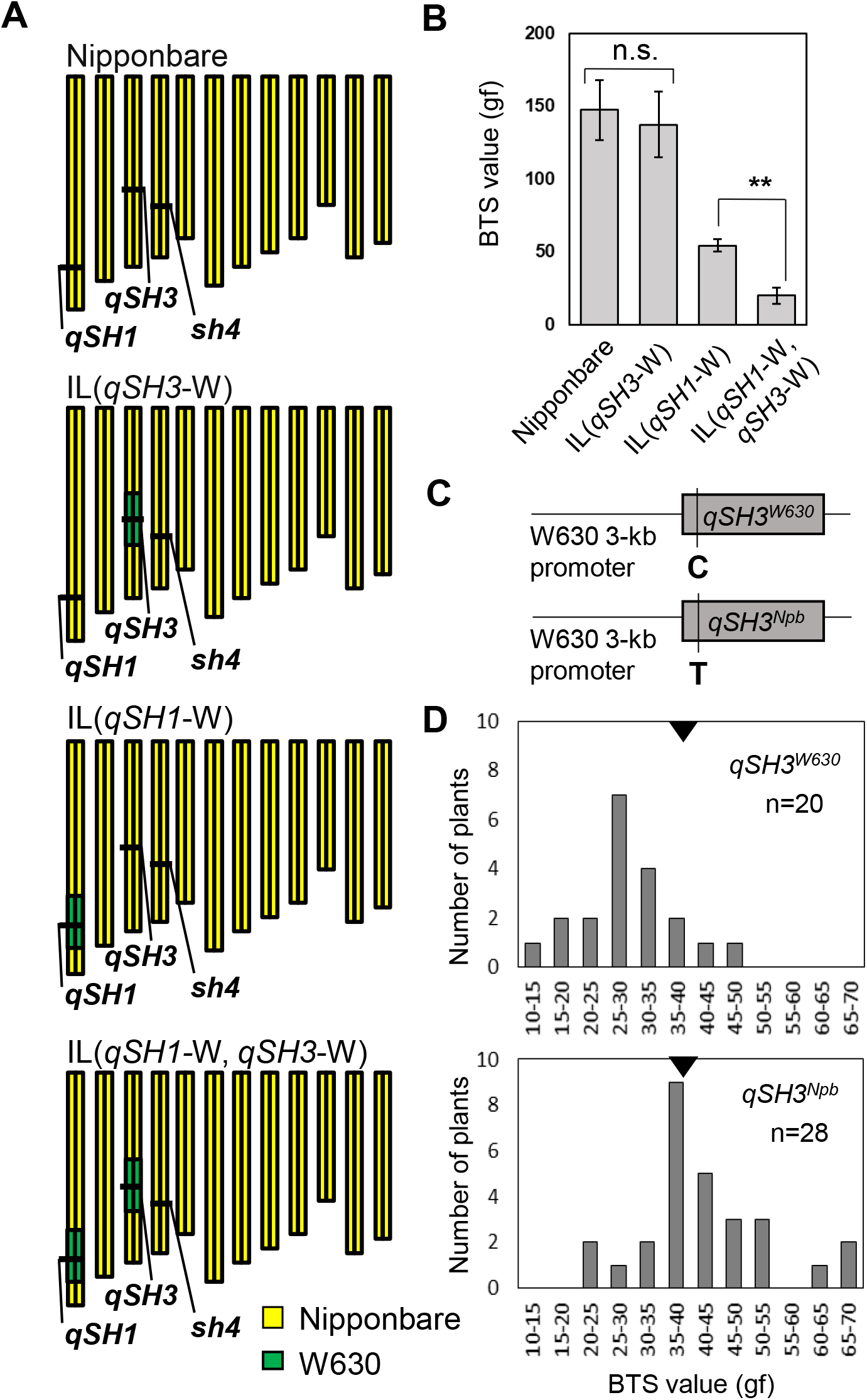
Identification of a causal SNP of *qSH3* associated with the degree of seed shattering. (*A*) Graphical genotypes of *O. sativa* Nipponbare and introgression lines (ILs) for *qSH3* and *qSH1* in the genetic background of cultivated rice, *O. sativa* Nipponbare. (*B*) Comparison of seed-shattering degree by breaking tensile strength (BTS) values in three ILs, IL(*qSH3*-W), IL(*qSH1*-W), and IL(*qSH1*-W, *qSH3*-W) in the Nipponbare genetic background. Data are mean ± S.D. of four plants. n.s. and ** indicate not significant and significant at the 1% level based on unpaired Student’s *t*-test, respectively. (*C*) Two types of constructs carrying *qSH3* cDNA sequences of W630 and Nipponbare (*qSH3*^*W630*^ and *qSH3*^*Npb*^) driven by a 3-kb region of the promoter used for transgenic analysis. (*D*) The BTS values observed for the transformants with *qSH3*^*W630*^ and *qSH3*^*Npb*^. Black triangles represent the average BTS value of IL(*qSH1*-W).

### Causal mutation at *qSH3* is conserved in both *japonica* and *indica* but not in *circum-*aus rice cultivars

Next, we analysed the distribution of SNP-70 at *qSH3* in cultivated rice. First, genotyping at the causal SNPs of *sh4, qSH1*, and *qSH3* was conducted for the three rice cultivars, Nipponbare, IR36, and Kasalath, belonging to *japonica, indica*, and *circum*-aus, respectively. All cultivars possessed the causal mutation at *sh4* and only Nipponbare had the causal mutation at *qSH1*. As for *qSH3*, Nipponbare and IR36 carried the causal mutation, but was absent in Kasalath (Fig. 2A). We further found that the Kasalath haplotype is similar to W630 around the *qSH3* region (Table S2). Using the diverse varieties of the World Rice Core Collection (WRC) (13), we found that 14 lines, all belonging to *circum*-aus, carried the functional allele at *qSH3* (Table S3 and Fig. S10 and S11). Furthermore, the Rice 3K genome project data (14) clearly showed that both *indica* and *japonica* carried the causal mutation, but almost 90% of *circum*-aus rice carried a functional allele at *qSH3* (Fig. 2B). To further understand the footprint of *qSH3* selection, we analysed nucleotide diversity across the *qSH3* genomic region. Using sequences of the Rice 3K genome collection (14), we detected a selective sweep at *qSH3* in both *indica* and *japonica* but not in the *circum*-aus linage (Fig. 2C, Fig. S12 and S13). These results suggest that *circum*-aus followed a separate trajectory to evolve reduced seed shattering (15). Thus, the reduction in seed shattering is dependent on lineage-specific variations in the subspecies, which is key to understanding the parallel processes of rice domestication.

**Fig. 2.**
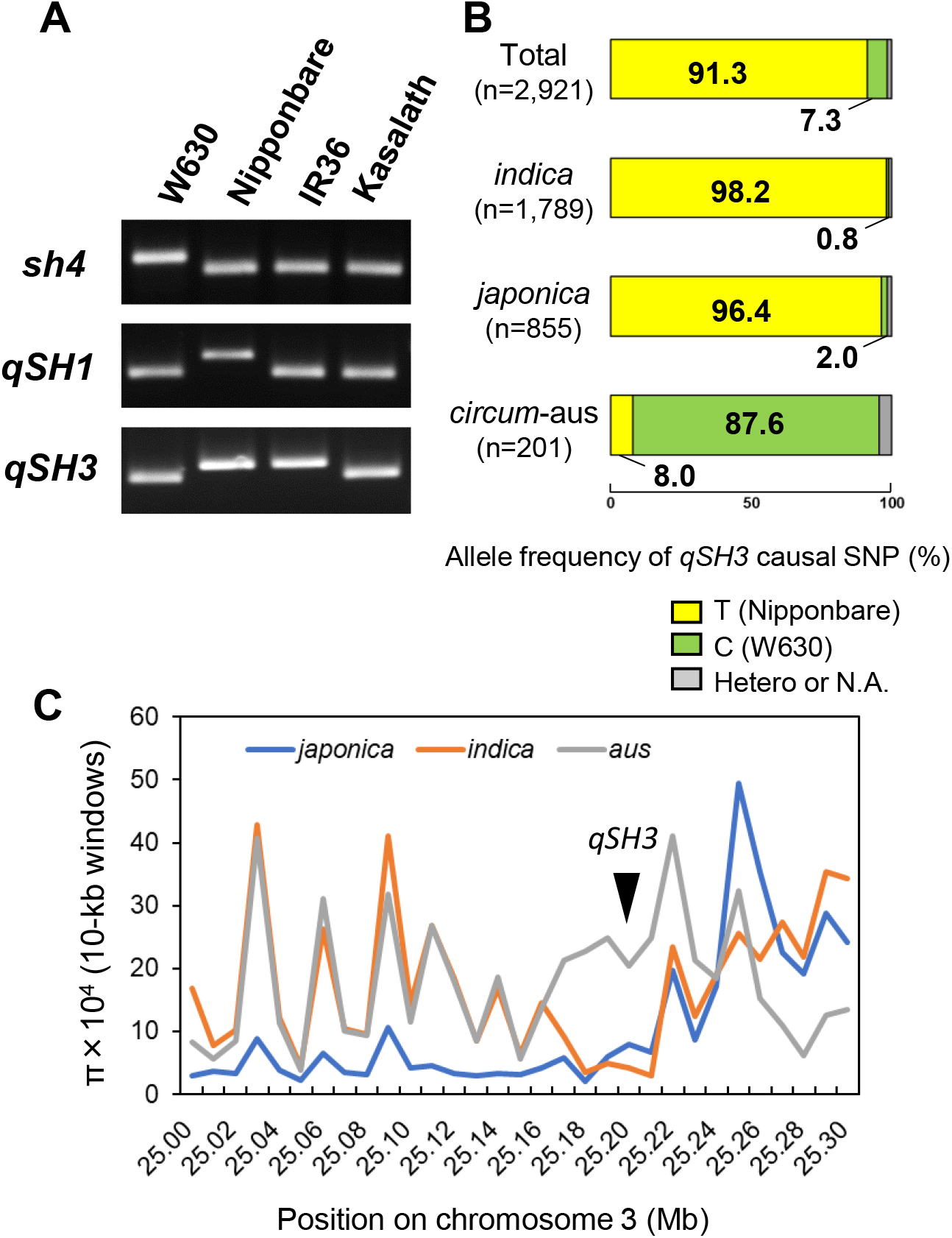
Lineage-specific selection at the *qSH3* locus in rice. (*A*) Genotyping at *sh4, qSH1*, and *qSH3* for *O. rufipogon* W630, *O. sativa japonica* Nipponbare, *indica* IR36, and *circum*-aus Kasalath based on the causal SNPs identified by dCAPS markers. (*B*) Allele frequency of *qSH3* causal SNP (%) in cultivated rice based on the 3K rice project data. Nipponbare type (‘T’) and W630 type (‘C’) are shown in yellow and green, respectively. (*C*) Nucleotide diversity (π) observed for domesticated rice in the physical position of 25.0–25.3 Mb on chromosome 3. In a flanking region of the *qSH3* locus (around 25.2 Mb), π was substantially decreased in *japonica* and *indica* cultivars. π was calculated in 10-kb windows using the SNP data of the 3K rice project (13).

### Role of *qSH3* causal mutation in an initial loss of seed shattering

We next aimed to understand how the causal SNP at *qSH3* contributed to initial rice domestication by reducing seed shattering in *japonica* and *indica*. Since the *sh4* mutation is conserved in all cultivated rice (Table S3), we produced IL(*sh4*-N) and IL(*qSH3*-N) in the genetic background of wild rice *O. rufipogon* W630. Evaluation of the seed-shattering mutations in the wild rice genetic background provides clearer morphological information related to the trait in early rice domestication. Complete formation of the abscission layer similar to that of W630 was observed in both ILs (Fig. 3), confirming that the single mutation at each locus was insufficient for phenotypic change in the abscission layer formation (7, 8). However, a slight inhibition of the abscission layer formation around vascular bundles was observed in IL(*sh4*-N, *qSH3*-N) (Fig. 3). Furthermore, a slight abscission layer inhibition was also observed in several wild rice accessions of *O. rufipogon* carrying mutations at both *sh4* and *qSH3* (Fig. S14 and S15, and Table S4), although they may have gained these domestic-type alleles through introgressions from cultivated rice (16). Even if the mutations in the two loci displayed a phenotypic difference in the abscission layer formation adequate to reduce shattering under greenhouse conditions without wind, their effect on seed shattering was much less than expected under field conditions.

**Fig. 3.**
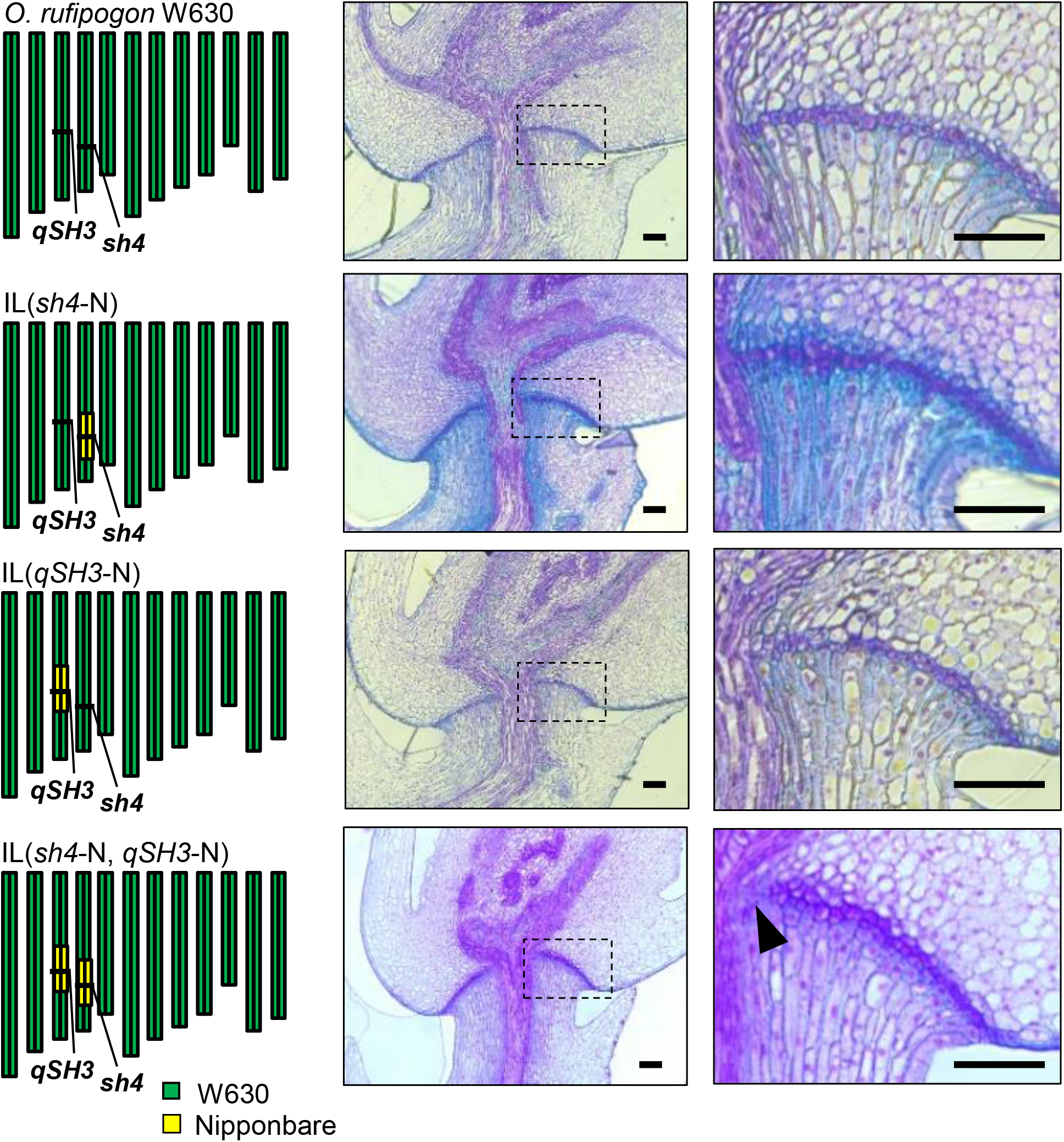
Abscission layer formation partially inhibited by *sh4* and *qSH3* in wild rice. Graphical genotypes of three introgression lines (ILs) in the genetic background of *O. rufipogon* W630 are shown on the left. Abscission layer formation in *O. rufipogon* W630, IL(*sh4*-N), IL(*qSH3*-N), and IL(*sh4*-N, *qSH3*-N) are shown. Each enlarged section indicated by the dotted square is shown in the righthand panel. A black arrowhead indicates an inhibited area of the abscission layer observed for IL(*sh4*-N, *qSH3*-N). Scale bars = 100 μm.

### Seed-shattering behaviour associated with a slight inhibition of abscission layer formation was mitigated by closed panicle formation

Previously, we reported that the closed panicle trait controlled by *OsLG1* had a major effect in rice domestication by facilitating grain harvest (7). A closed panicle reduces seed shedding by retaining seeds that get stuck by long awns found in the lower sections of the panicle. Interestingly, the *SPR3* (a locus regulating *OsLG1*) region is under strong selection in *indica, japonica*, and *circum*-aus (Fig. S16), suggesting that the closed panicle trait was selected in the early phase of rice domestication. Thus, closed panicles might have been associated with seed-shattering changes, although this trait, unlike seed shattering, is not visible archaeologically. Therefore, we generated seven wild ILs with different combinations of the three loci (*sh4, qSH3*, and *SPR3*) (Fig. S17) and compared their morphology together with that of the wild rice *O. rufipogon* W630 (Fig. S18). The heading date of all ILs was similar to that of W630 (Fig. S19); but there was a slight difference in inhibition of abscission layer formation around the vascular bundles depending on the double mutations at *sh4* and *qSH3* (Fig. 3 and Fig. S20), and open or closed panicle structure depending on *SPR3* mutation (Fig. 4A and 4B). Small BTS values were observed only for IL(*sh4*-N, *qSH3-*N) and IL(*sh4*-N, *qSH3-*N, *SPR3*-N) due to a slight inhibition of abscission layer formation, while the rest were close to zero as observed in wild rice (Fig. 4C). We were interested in assessing the differences in yields brought about by the causal mutations and identifying the combination of alleles that would have provided humans in prehistory with higher yields, including whether a closed panicle conferred additional value to humans when gathering wild rice. We therefore subjected the seven ILs and W630 to a seed-gathering experiment in the field (Fig. S21, Movies S1 and S2). The seed-gathering rates from three ILs with open panicle, namely IL(*sh4*-N), IL(*qSH3-*N), and IL(*sh4*-N, *qSH3*-N), were not significantly different from that of W630, regardless of the presence or absence of abscission layer inhibition (Fig. 4D and Table S5). However, the three ILs with a closed panicle structure and complete abscission layer formation, namely, IL(*SPR3*-N), IL(*sh4*-N, *SPR3*-N), and IL(*qSH3*-N, *SPR3*-N), presented a slightly increased yield (Fig. 4D). In contrast, IL(*sh4-*N, *qSH3*-N, *SPR3*-N), with a combination of closed panicle and a slight abscission layer inhibition, showed a significant increase in gathering rate compared with W630 (Fig. 4D and Table S5). As closed panicles would be easily visible in the field, humans could have targeted this higher yielding rice (Fig. S22), but even indiscriminate gathering also would retain larger proportions of grains from closed panicles.

**Fig. 4.**
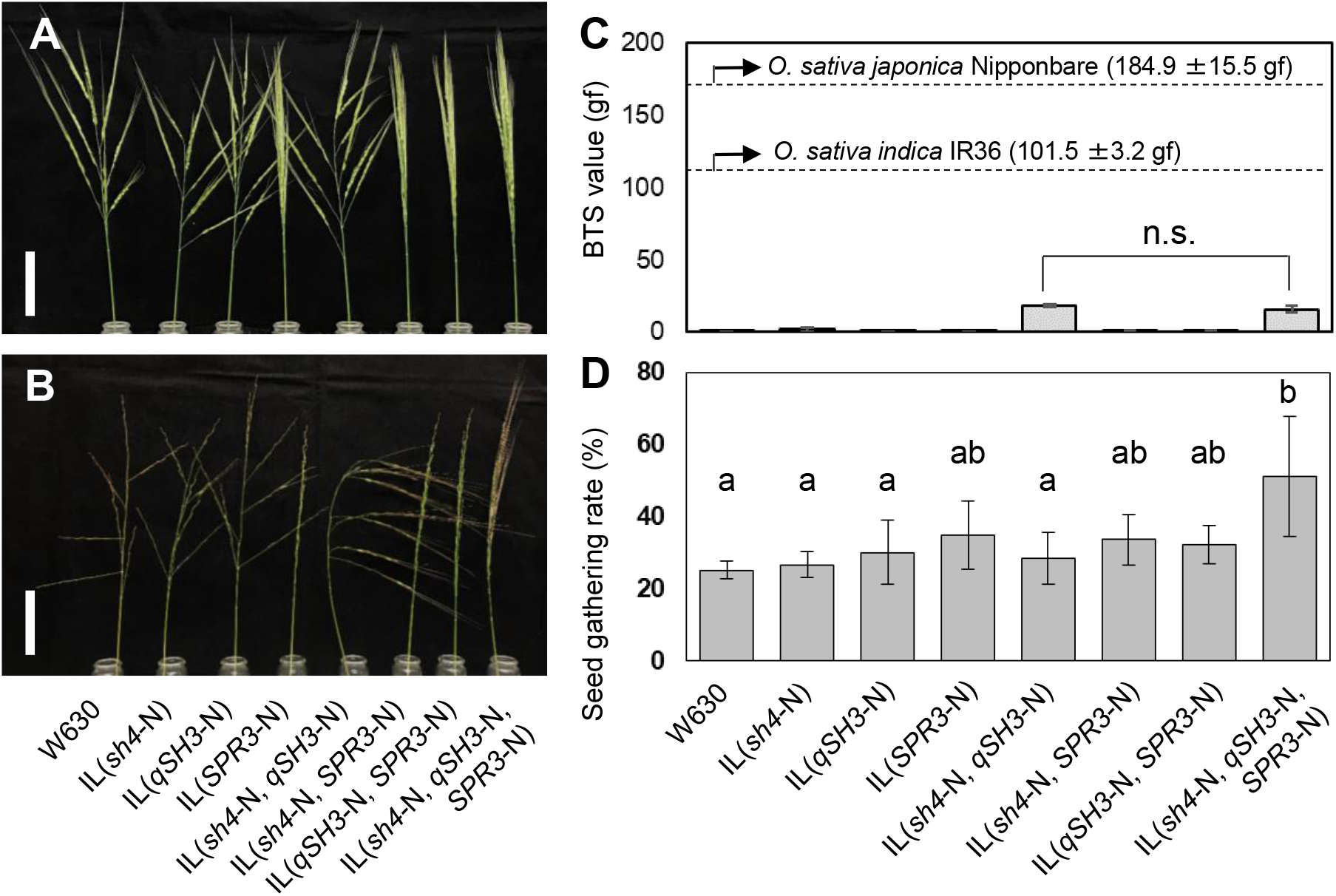
Role of *sh4, qSH3*, and *SPR3* mutations on the initial loss of seed shattering in rice domestication. (*A, B*) Panicle shape of the seven ILs at flowering (*A*) and seed-maturation (*B*) stages. Scale bars = 5 cm. (*C*) Breaking tensile strength (BTS) values for the seven ILs. BTS values for *O. sativa japonica* Nipponbare and *indica* IR36 are shown as controls. n.s. indicates not significant by unpaired Student’s *t*-test. (*D*) Seed-gathering rates (mean ± S.D. of four plot replicates, with nine plants in each plot) for *O. rufipogon* W630 and the seven ILs. Mean values labelled with different letters are significantly different, whereas those with the same letters are not (Tukey’s test with arcsine transformed values, *P* < 0.05).

When plants were sown from harvests, closed panicles and reduced seed shattering would increase, owing to the enhanced gathering rate that would occur in the presence of all three the domesticated-type alleles. As these alleles increased in frequency and became fixed, yields would have increased, which would have encouraged further investment in rice cultivation (17). The selection for closed panicles instigates self-pollination behaviour owing to the long awns, which disturb the free exposure of anthers and stigmas (stigmas and closed panicle) (7). Thus, a closed panicle may also be advantageous in mitigating natural variations in seed-shattering loci by reducing outcrossing. Although awns are undesirable in modern cultivated rice, gathering of rice in the early stages of rice domestication might have benefitted from the presence of awns in plants with closed panicles, as these would have increased yield and self-pollination rates.

### Complementary interaction of a slight inhibition of abscission layer and closed panicle formation synergistically contributed to structural stability of the panicle increasing yield

To better understand how the closed panicle, caused by *SPR3*, and the inhibition of abscission layer formation, caused by *sh4* and *qSH3*, contributed to the initial loss of seed shattering, we analysed their roles by performing structural mechanics analysis. The awns of wild rice play a pivotal role in seed dispersal in spreading panicles (Fig. 5A). We measured the lengths and weights of awns and grains in wild rice *O. rufipogon* W630 (Fig. 5B), and using these values, we calculated the sectional force exerted on the spikelet base depending on panicle angles (Fig. S23). The axial and shear forces were slightly increased in a closed panicle compared to an open panicle. However, the bending moment, which is the predominant factor affecting seed dispersal in an open panicle, was considerably reduced in a closed panicle (Fig. S23). A slight inhibition of the abscission layer formation by *sh4* and *qSH3* led to an increase in the length of abscission layer inhibition (Fig. 3). Therefore, we measured the length of the abscission layer and central vascular bundle in *O. rufipogon* W630 by scanning microscopy (Fig. 5C and 5D). We also calculated the moment of inertia of the area, a property that describes the torque required to break the abscission layer and so disarticulate the grain, for the disrupted abscission layer (Fig. S23). The value increased exponentially with increasing lengths of abscission layer inhibition. A reduction in the bending moment and an increase in the moment of inertia of the area act synergistically to reduce bending stress (Fig. 5E), contributing to the structural stability of spikelets without shattering. Thus, the interaction between a closed panicle and abscission layer inhibition acted complementarily to increase yield.

**Fig. 5.**
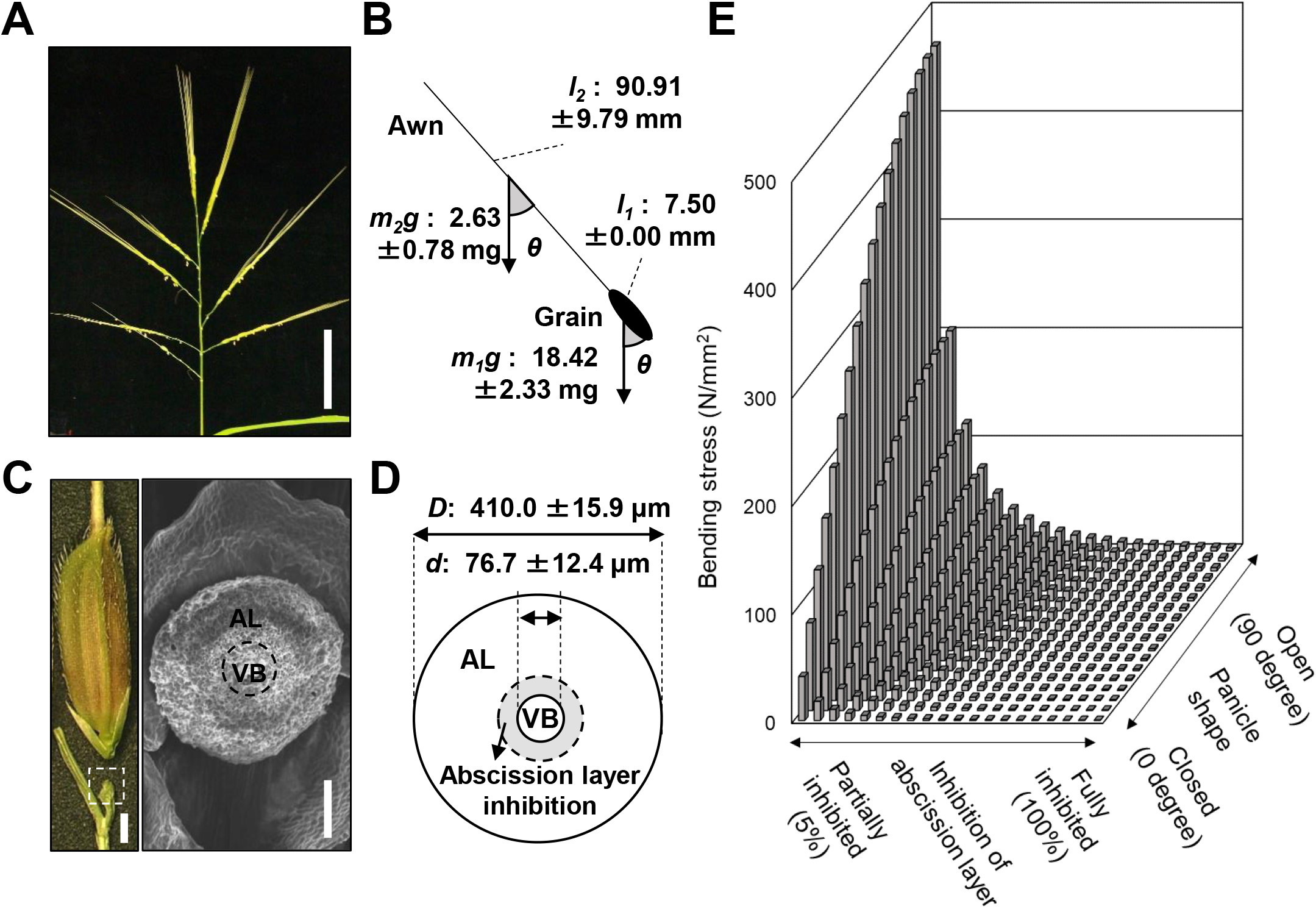
Structural mechanics analysis of panicle shape and abscission layer inhibition related to the initial loss of seed shattering. (*A*) Panicle shape of *O. rufipogon* W630. Bar = 5cm. (*B*) Schematic representation of awn and grain with panicle angle. *θ* represents the panicle angle. *l*_*1*_, *l*_*2*_, *m*_*1*_*g*, and *m*_*2*_*g* indicate the length of grain, length of awn, weight of grain, and weight of awn, respectively. (*C*) Detachment of grain and pedicel (left), and scanning electron microscopy (SEM) analysis of the pedicel abscission layer of *O. rufipogon* W630 (right). AL and VB indicate abscission layer and vascular bundle, respectively. Bars = 1 mm (left) and 100 μm (right). (*D*) Schematic representation of the abscission layer. *D* and *d* indicate the diameter of the abscission layer and vascular bundles, respectively. Dotted circle indicates the area of disrupted abscission layer. (*E*) Simulation of the bending stress exerted on the spikelet base depending on panicle shape and abscission layer inhibition. A higher stress was experienced in the open panicle with less inhibited abscission.

## Conclusions

In this study, we identified key functional genes that contributed to rice domestication by increasing harvest yields. We identified the causal SNP at *qSH3* involved in reduced seed shattering, and demonstrated that the previously proposed *sh4* mutation alone could not trigger the non-shattering morphology in rice (6, 7, 16). We also explained how the early selection of a closed panicle and the resulting mechanics of spikelet retention would have increased yields and facilitated the selection for non-shattering. Our results showed that the initial step in rice domestication was more complex than previously thought. Based on information from our studies, changes in the long-held perspective from a single domestication allele model to one where synergistic effects of several domestication genes are needed.

The origin and spread of rice subspecies based on population genetic analysis have been the subject of discussions (15, 18–21). However, most of these studies have been conducted using the genome information of modern cultivars and wild rice, without considering the importance of visible phenotypic changes that could have been targeted by humans. Therefore, the lineage-specific variations associated with quantitative traits are a key to further understanding the process of rice domestication.

Based on our work, we propose a stepwise route for rice domestication. In wild rice, any of the natural variations in the loci for seed shattering and closed panicle formation alone had little effect on increasing yield. Their combination, however, established an archaic rice which could have been visibly recognised by ancient gatherers as advantageous to increasing yields. The change in harvesting efficiency along with the use of harvesting tools further promoted the selections of natural variants in domestication-related traits in rice, a crop which now supports billions of people worldwide.

## Materials and Methods

Details regarding plant materials, QTL mapping, fine mapping, transformation, population genetic analysis, histological analysis, seed gathering experiment, and structural mechanics analysis are provided in the Supplementary Information. Primers used for genetic mapping, gene expression analysis, production of transgenic plants, and genotyping are shown in Table S6-S8.

## Acknowledgements

The wild rice accessions of *O. rufipogon* were provided by the National Institute of Genetics supported by the National Bioresource Project, MEXT, Japan. We thank R. Morita and H. Fukayama, Kobe University, for their support in producing transgenic plants, and H. Furuumi, National Institute of Genetics, for growing wild rice accessions. This work was partly supported by Grants-in-Aid from the Japan Society for the Promotion of Science, 15KK0280 and 18K05594 (R.I.), JSPS overseas research program (C.C.C.), JSPS Bilateral Open Partnership Joint Research Projects, Nos. JPJSBP120189948 and JPJSBP120219922 (R.I., D.F.), Nikki Saneyoshi and Kinoshita foundations (R.I.), and NIG-Joint research (K-I.N, T.I.).

