## Supplementary Information for "A stepwise route to domesticate rice by controlling seed shattering and panicle shape"

### Materials and Methods

#### Plant materials and growth conditions

Wild rice accession of *Oryza rufipogon* W630 originated from Myanmar, cultivated rice *O. sativa japonica* Nipponbare, *indica* IR36, and *circum*-aus Kasalath were used in this study. *O. rufipogon* accessions provided by the National Institute of Genetics were also used.

#### Evaluation of seed-shattering degree

The degree of seed shattering was evaluated by measuring the breaking tensile strength (BTS, gf: gram force) with a digital force gauge (FGP 0.5, Nidec-Shimpo, Japan). This value is the required amount of tensile stress needed to detach a grain from the pedicel. The BTS values of seeds (25 randomly selected seeds from three panicles) were measured, approximately a month after heading, and the average BTS values were calculated.

#### Histological analysis of abscission layer formation

Tissue samples of the pedicel of grains were collected for histological analysis before heading, as previously reported (1, 2). The samples were cut into 3-μm sections using a rotary microtome, RM1215RT (Leica Biosystems, Germany), and then stained with toluidine blue O solution. These sections were photographed with a digital camera using the imaging software, ToupView ( × 86) (Amscope.com, USA).

#### Fine mapping of *qSH3*

To fine-map *qSH3*, plants were screened for those with the recombination between RM3513 and RM3601, and eleven recombinants were obtained. Further progeny test showed that the candidate region was found to be between RM15584 and RM15586, a 41-kb region (Fig. S2). The region harboured a part of single gene, *Os03g0650000*, encoding a rice YABBY transcription factor (Fig. S2). Two critical recombinant plants, nos. 108-15-97-4 and 108-15-97-27 were further analysed for their recombination positions between RM15584 and *qSH3*-E2, and *qSH3*-E2 and RM15586, respectively, using polymorphic markers (Table S6). A sequence survey in the 1,228 bp region of the 5' UTR, exon 1, and intron 1 of *qSH3* identified seven polymorphisms between Nipponbare and W630 (Fig. S4 and Table S2). Of these, one SNP, namely SNP-70, was found in the exon region of *OsSh1*, that caused amino acid change of leucine (W630) and phenylalanine (Nipponbare) (L24F).

**Production of Introgression Lines (ILs) for *qSH1* and *qSH3* in Nipponbare background**

To analyse the effects of *qSH3* on seed shattering, ILs in the genetic background of Nipponbare were produced. Backcross recombinant inbred lines (BRILs) in a cross between Nipponbare, as recurrent parent, and W630, as donor parent were previously developed (3). Two BRILs, namely AsN141 and AsN175, were selected based on the genotypes at *qSH1* and *qSH3*. These BRILs had W630 chromosomal segments on chromosomes 1 and 2 for AsN141 and on 3, 6, and 11 for AsN175. The two BRILs were then crossed and resulting $F_2$ and $F_3$ plants were surveyed for the genotypes at the two loci (*qSH1* and *qSH3*) with W630 chromosomal segment(s). The other regions were fixed with Nipponbare chromosomal segments. Furthermore, three ILs, IL(*qSH1*-W), IL(*qSH3*-W), and IL(*qSH1*-W, *qSH3*-W), were produced as well (Fig. 1A). The IL(*qSH1*-W) was used for the transformation test.

**Complementation test for *qSH3* causal SNP by transgenic rice**

To evaluate if the SNP at *qSH3* is associated with seed-shattering degree, a transformation experiment was performed. Two types of constructs having the 3-kb W630 promoter region of *qSH3* (*OsSh1*) and cDNA with Nipponbare (T) or W630 (C) alleles at the *qSH3* causal SNP were generated (Fig. S7). These were transformed into IL(*qSH1*-W). The 3-kb promoter region of *qSH3* (*OsSh1*) was amplified from genomic DNA of W630 using KOD-FX-plus polymerase (TOYOBO, Japan) and a pair of primers with restriction sites (*Xho*I/*Spe*I and *Sma*I) (Table S7). For the plasmid construction of *qSH3* transgene, the amplified cDNA was first prepared using total RNA extracted from W630 and Nipponbare leaves with primer having restriction enzyme sequence, *Sma*I and *Sac*II on the reverse primer (Table S7). These vectors called $qSH3^{W630}$ and $qSH3^{Npb}$ contained W630 *qSH3* promoter region and the candidate causal SNP at *qSH3* of W630 (C) or Nipponbare (T) type nucleotide. The resulting constructs ($qSH3^{W630}$ and $qSH3^{Npb}$) in pBI101-Hm3 were introduced into IL(*qSH1*-W) using *Agrobacterium tumefaciens* (EHA105)-mediated transformation. After screening of the transgenic line based on the expression analysis for the *qSH3* transgenes (Table S8), transformants were evaluated for their breaking tensile strength values from 25 randomly selected seeds from a panicle using a digital force gauge (FGP 0.5, Nidec-Shimpo, Japan).

**RNA extraction and expression analysis of *qSH3***

To compare *qSH3* gene expression levels between Nipponbare and W630, total RNA was extracted from spikelet base tissue before heading. To screen transgenic plants expressing $qSH3^{W630}$ or $qSH3^{Npb}$ transgene, total RNA was extracted from leaves in order to save the spikelet tissue for evaluation of seed-shattering degree. RNA extraction was carried out using Plant Total RNA Mini Kit (VIOGENE, Taiwan). After a DNase I treatment, cDNA was prepared using oligo dT primer and ReverTra Ace qPCR RT Master Mix with gDNA Remover according to the manufacturer's protocol (TOYOBO, Japan). The RT-PCR was carried out using *Quick Taq* polymerase (TOYOBO, Japan) according to the manufacturer's protocol with gene-specific primer sets (Table S7 and S8). Expression levels of *qSH3* between Nipponbare and W630 was measured by quantitative real-time PCR (qPCR) using THUNDERBIRD® SYBR qPCR Mix (TOYOBO, Japan) with a Light Cycler 96 (Roche, Tokyo, Japan) and the gene-specific primer sets (Table S7). Actin was used as an internal control.

**Distribution of the causal SNP at *qSH3* and the linked SNP at *SPR3* in cultivated rice**

We genotyped three seed-shattering loci, *sh4*, *qSH1*, and *qSH3*, for *O. rufipogon* W630, *O. sativa* Nipponbare, IR36, and Kasalath using dCAPS markers specifically detecting the causal SNPs (Table S8). We also genotyped *qSH3* allele of SNP-70 in the world rice core collection (WRC) (4). The genotypes of the SNP-70 were also confirmed with whole-genome sequencing data publically available from the TASUKE system, a web browser visualization system for whole-genome variant data (5). The genotypes of *qSH3* causal SNP (SNP-70) for *japonica*, *indica*, and *circum*-aus rice cultivars were also investigated using Rice SNP-Seek Database (6, 7). In addition, although the causal mutation at *SPR3* is not known (8), genotype of the SNP linked to *SPR3* (9) was investigated in a similar manner.

**Nucleotide diversity analysis of *qSH3* and *SPR3***

Estimation of nucleotide diversity was based on "3K RG 29mio biallelic SNPs Dataset" (Rice SNP-Seek Database: https://snp-seek.irri.org/_download.zul). We selected 772 individuals of *japonica* (tropical, sub-tropical, and temperate), 1,174 individuals of *indica* (ind1A, 1B, 2, and 3), and 201 individuals of *circum*-aus as "non-admixed" individuals which have not undergone introgression from other subpopulations. Loci with >50% missing genotypes were removed with PLINK (10) and haplotype phasing was performed for chromosomes 3 and 4 using Beagle 5.0 (11). Then, we removed loci with minor allele frequency (MAF) <0.02 and calculated nucleotide diversity (π) in 10-kb non-overlapping sliding windows for each subpopulation using selscan (12). For an estimation of regions that experienced positive selection on chromosome 3, we carried out the extended haplotype homozygosity (EHH)-based methods, nSL (13) for single populations and XP-EHH (14) for dual population comparisons with selscan. Unstandardized nSL and XP-EHH values for each core SNP were Z-scored and transformed to p-values.

**Evaluation of *sh4* and *qSH3* genotypes at their causal SNPs in wild rice accessions**

Genotypes of *sh4* and *qSH3* in *O. rufipogon* accessions were investigated using OryzaGenome (http://viewer.shigen.info/oryzagenome2detail/index.xhtml) (Fig. S14). The *sh4* genotype of wild rice was unavailable in OryzaGenome. From 446 accessions, 80 were found with the Nipponbare type cultivated allele at the *qSH3* causal SNP (SNP-70). Of these, 49 accessions were available for distribution from the National Institute of Genetics in Mishima, Japan. Genotyping of these 49 accessions at *sh4* was carried out using dCAPS marker (Table S8), and identified the 27 accessions were identified to carry the Nipponbare-type cultivated allele at *sh4*. These plants were grown at Kobe university or National Institute of Genetics, where short day treatment was applied to induce their flowering. A total of 17 accessions were available following flowering and were evaluated for their abscission layer formation.

**Production of ILs for *sh4*, *qSH3*, and *SPR3* in *O. rufipogon* W630 background**

To evaluate the effect of *sh4*, *qSH3*, and *SPR3* on initial loss of seed shattering, ILs of the combination at the three seed-shattering loci in the genetic background of wild rice, *O. rufipogon* W630, were produced. Two ILs named IL(*sh4*-N) and IL(*SPR3*-N) were previously developed by backcrossing in the genetic background of *O. rufipogon* W630. Similarly, IL(*qSH3*-N) was produced. IL(*sh4*-N, *qSH3*-N), IL(*sh4*-N, *SPR3*-N), IL(*qSH3*-N, *SPR3*-N), IL(*sh4*-N, *qSH3*-N, *SPR3*-N) were produced by crossing among IL(*sh4*-N), IL(*qSH3*-N), and IL(*SPR3*-N). Genotypes at the three loci were confirmed with dCAPS markers for *sh4* and *qSH3* (Table S8), and two flanking SSR markers for *SPR3* (8), respectively. Their chromosomal constitutions of these lines are shown (Fig. S17).

**Seed gathering experiment**

We estimated the seed-gathering rate by beating panicles to collect mature seeds (Movie S1 and S2). Seedlings of a wild rice accession, *O. rufipogon* W630, and the seven ILs were first geminated in the laboratory and then transplanted to a paddy field at Kobe University, Japan (34°43′ N, 135°14′ E) on 8 June 2017. A 40-cm square block with nine plants in rows of 3 plants × 3 plants were planted at intervals of 20 cm and were set as one experimental plot with four replicates (Fig. S21). Heading took place between 13-18 August 2017. To determine the seed setting ratio, ten panicles per plot were bagged on the 22-25 August 2017. They were collected on 1 September 2017 and the average seed setting ratios were evaluated (8). Regarding the number of seeds per panicle, 20 to 24 panicles per plot were further counted *in situ* in the field on 25 August 2017. The average numbers were calculated together with the ten bagged panicles. We gathered the seeds by beating panicles to collect mature seeds into a plastic pan. Harvesting by beating took place every 2 days with a start date on 28 August 2017 and an end date on 11 September 2017, a total of 8 times. The average time it took to harvest mature spikelets per plot was 2.5 minutes and it took approximately 2.5 to 3.0 hours to complete the entire field.

The total number of seeds produced in each plot was estimated using the following parameters (8): collected seed weight (CSW; total weight of fully filled seeds), 1,000-seed weight (1000SW; weight of 1,000 fully filled seeds collected), seed setting rate (SSR; average of ten panicles), number of seeds per panicle (NSP; average of 30-34 panicles), number of panicles (NP; total number of all effective panicles), number of seeds collected ((CSW/1000SW) × 1,000), total number of seeds produced (SSR × NSP × NP). Estimation of the seed-gathering rate was based on the percentage of the number of seeds collected against the total number of seeds produced. The average in seed-gathering ratio was calculated by Tukey's test to determine the significance of differences in seed-gathering rates among wild and the seven ILs and this is reported in the manuscript and Fig. 4.

**Structural mechanics analysis on panicle shape and abscission layer inhibition.**

To evaluate the effect of the interaction of a closed panicle formation by *SPR3* and slight inhibition of abscission layer formation by *sh4* and *qSH3*, structural mechanics analysis was performed. To calculate the sectional force exerted on the spikelet base, the length and weights of awns and grains of wild rice *O. rufipogon* W630 were measured. The average length of grain ($l_1$) and awn ($l_2$) were 7.50±0.00 and 90.91±9.79 mm, respectively, and the average weight of grain ($m_1g$) and awn ($m_2g$) were 18.42±2.33 and 2.63±0.78 mg, respectively. The angle of awn and grain against vertical axis was provided by θ, which changes depending on panicle structure (open to closed). Then, axial force (*NA*), shear force (*SA*), and bending moment (*MA*) were calculated (Fig. S23).

Axial force (*NA*) was calculated as:

$$NA = m_1 g \cos\theta + m_2 g \cos\theta \quad (1)$$

Shear force (*SA*) was calculated as:

$$SA = -m_1 g \sin\theta - m_2 g \sin\theta \quad (2)$$

Bending moment (*MA*) was calculated as:

$$MA = m_1 g \sin\theta \frac{l_1}{2} + m_2 g \sin\theta (l_1 + \frac{l_2}{2}) \quad (3)$$

Slight inhibition of the abscission layer formation by *sh4* and *qSH3* led to an increase in the length of abscission layer inhibition (Fig. 3). The vascular bundles located in the centre of the pedicel are vacant spaces without a cell connection. The inhibition of abscission layer formation by *sh4* and *qSH3* provided cells in the toroidal region that connecting pedicel and grain around vascular bundles, this is shaped like a circular hollow section. Then, the length of the diameter of the outer edges of abscission layer (*D*) and central vascular bundle (*d*) in *O. rufipogon* W630 were measured using scanning electron microscopy (Fig. 5C), as previously described (2). Then, the moment of inertia of the area, a property that predicts bending stress, was calculated.

The moment of inertia of the area (*I*) for the circular hollow section is calculated as:

$$I = \frac{\pi(D^4 - d^4)}{64} \quad (4)$$

The *D* value changes depending on the degree of inhibition of abscission layer formation. $D^{max}$ and $D^{min}$ are fully inhibited and completely formed abscission layer, respectively. Thus, $D^{max}$ and $D^{min}$ are equivalent values for completely non-shattering *japonica* rice cultivars and completely shattering wild rice, respectively. The average $D^{max}$ and $D^{min}$ (same as *d*) are 410.0 ±15.9 and 76.7 ±12.4 μm (n = 5, data are means ± S.D.), respectively.

Then the modulus section (*Z*) is calculated as:

$$Z = \frac{\pi(D^4 - d^4)}{32D} = \frac{2I}{D} \quad (5)$$

To understand the force exerted on the spikelet base inducing seed shattering, bending stress ($\sigma$) is calculated as:

$$\sigma = \frac{MA}{Z} \quad (6)$$

Reduction in bending moment (*MA*) associated with closed panicle formation by *SPR3* and increase in section modulus (*Z*) caused by an inhibition of abscission layer formation by *sh4* and *qSH3* drastically reduced bending stress. These produced the stable structure supressing seed shattering.

**Legends for Movies S1 and S2**

Movie S1. Growth conditions of wild rice *O. rufipogon* W630 and seven ILs used for harvesting experiment in a paddy field.

Movie S2. Example of seed gathering experiment.
